# Quantitative and rapid *Plasmodium falciparum* malaria diagnosis and artemisinin-resistance detection using a CMOS Lab-on-Chip platform

**DOI:** 10.1101/638221

**Authors:** K. Malpartida-Cardenas, N. Miscourides, J. Rodriguez-Manzano, L. S. Yu, J. Baum, P. Georgiou

**Affiliations:** Centre for Bio-Inspired Technology, Department of Electrical and Electronic Engineering, Imperial College London, UK; Department of Life Sciences, Imperial College London, UK

## Abstract

Early and accurate diagnosis of malaria and drug-resistance is essential to effective disease management. Available rapid malaria diagnostic tests present limitations in analytical sensitivity, drug-resistant testing and/or quantification. Conversely, diagnostic methods based on nucleic acid amplification stepped forwards owing to their high sensitivity, specificity and robustness. Nevertheless, these methods commonly rely on optical measurements and complex instrumentation which limit their applicability in resource-poor, point-of-care settings. This paper reports the specific, quantitative and fully-electronic detection of *Plasmodium falciparum*, the predominant malaria-causing parasite worldwide, using a Lab-on-Chip platform developed in-house. Furthermore, we demonstrate on-chip detection of C580Y, the most prevalent single-nucleotide polymorphism associated to artemisinin-resistant malaria. Real-time non-optical DNA sensing is facilitated using Ion-Sensitive Field-Effect Transistors, fabricated in unmodified complementary metal-oxide-semiconductor technology, coupled with loop-mediated isothermal amplification. This work holds significant potential for the development of a fully portable and quantitative malaria diagnostic that can be used as a rapid point-of-care test.

## 1 Introduction

Rapid diagnosis of infectious diseases at the point-of-care (PoC) is essential to enable effective surveillance of cases, infection control and appropriate treatment administration [1]. Lab-on-chip (LoC) diagnostic platforms have experienced significant growth in recent years as the result of huge advances in several disciplines such as biosensing technologies, molecular biology and microfluidics. Their potential use in resource-limited settings while providing clinical sensitivity, specificity, high speed of detection and an easy-to-use interface are key factors for epidemiological reporting of antimicrobial-resistance and disease management [2].

Various techniques have been employed to identify the presence of pathogens in infected patients, targeting either the pathogen itself, biological products derived from the pathogen, or alterations in patients’ biomolecules such as protein/nucleic acid overexpression. These techniques can be classified into three main categories: (i) cellular-based methods, (ii) protein-based methods and (iii) nucleic acid-based methods. Specifically, cellular-based methods such as microscopy, are commonly used for pathogen identification, but require high expertise and expensive equipment thus limited to centralized facilities [3–7]. Conversely, most reported protein-based methods rely on antigen-antibody detection and are typically combined with paper-based diagnostics such as lateral flow assays (LFAs) [8–10]. Even though they report results within 10 to 20 min, they commonly suffer from low sensitivity or low specificity for clinical applications. In addition, they mainly rely on colorimetric or fluorescence measurements which are not usually capable of quantification unless expensive equipment such as optical cameras or lasers are involved [11]. Instead, nucleic-acid amplification tests (NAATs) are characterized by high sensitivity and specificity, enabling quantification and detection of early stage infections. Among them, polymerase chain reaction (PCR) is the most widely used technique and is currently considered the gold standard in centralized laboratories. However, the requirements for thermal cycling and expensive equipment limit its application in PoC diagnostics. Isothermal methods have emerged as the next generation of NAATs, due to their capability of running at constant temperature which reduces equipment complexity. In particular, loop-mediated isothermal amplification (LAMP) has received considerable attention due to its increased detection speed compared to PCR (less than 35-40 min), higher sensitivity as well as higher specificity owing to the higher number of primers used [12]. These features make LAMP an attractive NAAT method for the PoC diagnosis of infectious diseases [13–15].

One of the most threatening infectious diseases in resource-limited settings, malaria, with an estimated 219 million cases in 2017, has been identified by the World Health Organization (WHO) as one of the main targets to be addressed [16]. Malaria is caused by the *Plasmodium* parasite, with five species identified to infect human population out of which *Plasmodium falciparum* is the most prevalent. In addition, the emergence of drug-resistant *P. falciparum* strains to medicines such as artemisinin, is compromising the effectiveness of current malaria treatments. Consequently, there is a significant need to specifically detect this pathogen with high sensitivity and rapid detection for adequate therapy prescription. Most rapid diagnostic tests developed for this purpose are no longer reliable due to the loss of expression of targeted proteins, such as HRP-II (leading to high occurrence of false negatives) [17, 18]. As a result, subsequent efforts have focused their attention on NAATs, primarily targeting the gene *18S rRNA*. Nevertheless, significant challenges towards rapid diagnosis sill exist with (i) gold standards being PCR-based which rely on thermal cycling and reported time-to-results longer than 1 hr, (ii) most of the reported LAMP primer sets [19–21] did not consider the zoonotic *Plasmodium* species *P. knowlesi* which has recently jumped to infect human hosts [22, 23], (iii) most of the reported NAATs are lab-based and non-portable and (iv) if targeting PoC applications, they rely on optical measurements which typically only provide qualitative results. A summary of the most relevant NAATs for *P. falciparum* identification is provided in Table 1.

**Table 1.** Nucleic-acid amplification tests for the specific detection of *P. falciparum*.

| Assay | Target (gene) | Limit of Detection | Time to Positive | Species considered | Ref. |
| --- | --- | --- | --- | --- | --- |
| Nested PCR | 18S rRNA | 6 p/μL<sup>a</sup> | > 60 min | Pv, Po, Pm* | Singh *et al.* [24] |
| Nested PCR | K13 | NA<sup>b</sup> | > 60 min | Pv, Po, Pm, Pk | Talundzic *et al.* [25] |
| PCR | 18S rRNA | 1 c/r | > 60 min | Pv, Po, Pm | Mangold *et al.* [26] |
| LAMP | 18S rRNA | 100 c/r<sup>c</sup> | 32 min | Pv, Po, Pm | Han *et al.* [21] |
| LAMP | 18S rRNA | 5 p/μL | 35 min | Pv, Pm* | Mohon *et al.* [19] |
| LAMP | 18S rRNA | 100 p/μL | > 60 min | Pv, Po, Pm* | Poon *et al.* [27] |
| LAMP | 18S rRNA | 1 c/r | 35 min | Pv, Po, Pm, Pk* | Lau *et al.* [20] |
| LAMP | mitochondrial<sup>d</sup> | 5 p/μL | 30-40 min | Pv, Po, Pm, Pk | Polley *et al.* [28] |
| LAMP<sup>e</sup> | ND<sup>f</sup> | 1 p/μL | 45 min | ND<sup>f</sup> | Loopamp Kit [29] |
| LAMP<sup>g</sup> | K13 | 1 c/r | < 20 min | Pv, Po, Pm, Pk | **This work** |
| pH-LAMP<sup>g</sup> | K13 | 10 c/r | < 25 min | Pv, Po, Pm, Pk | **This work** |

\* *Plasmodium* species mentioned in the corresponding manuscript, with no cross-reactivity results among species shown.
<sup>a</sup> Parasites per μL of blood (p/μL). <sup>b</sup> Not Available. <sup>c</sup> Copies per reaction (c/r). <sup>d</sup> DNA. <sup>e</sup> Qualitative.
<sup>f</sup> Non Disclosed. <sup>g</sup> Quantitative.
This table displays information as stated in company websites, product specification sheet, or references and does not intend to cover all nucleic-acid diagnostic tests for *P. falciparum* detection.

As opposed to optical-based detection, electrochemical detection is more promising due to innate capabilities such as high sensitivity, large scale integration, detection on miniaturized hardware, fast response and quantifiable measurements [30, 31]. Field-effect transistors and more specifically ion-sensitive field-effect transistors (ISFETs) are emerging potentiometric sensors for NAAT applications. Owing to their compatibility with CMOS technology (complementary metal-oxide-semiconductor), fully-electronic chemical detection is possible while ensuring sensor miniaturisation, mass manufacturing and low cost. CMOS-based ISFET sensing has been demonstrated for the detection of DNA amplification by employing an adapted version of LAMP, called pH-LAMP, that allows changes in pH to occur during nucleic acid amplification such that they can be detected by the ISFET sensors [32–34]. Furthermore, this technology is a suitable candidate for point-of-care implementations through transferring the amplification chemistries on a Lab-on-Chip platform with integrated sensing.

In this paper, we report the rapid and specific detection of *P. falciparum* malaria using a novel molecular assay targeting the gene *kelch 13*. Detection takes place in less than 20 minutes in isothermal conditions using LAMP. Furthermore, we show pH-LAMP based detection of malaria on a Lab-on-Chip platform which uses ISFETs to facilitate direct chemical-to-electronic sensing. The LoC platform demonstrates for the first time DNA quantification on-chip, using DNA samples derived from clinical isolates of *P. falciparum*. In addition, we further demonstrate on the LoC platform, detection of the *C580Y* single-nucleotide polymorphism (SNP) associated with artemisinin-resistant malaria. Overall, the proposed molecular methods in combination with ISFET-based sensing are capable of label-free amplification and quantification of nucleic acids and thus lend themselves to PoC implementations of any desired target (including both DNA and RNA) with high sensitivity, specificity and speed of detection.

## 2 Results

### 2.1 Analytical specificity of the *P. falciparum* LAMP primer set

A region of the kelch propeller domain within the gene *kelch 13* was selected as the target for *P. falciparum* specific detection, as shown in Fig. 1A. In contrast to the commonly reported gene 18S rRNA, the gene *kelch 13* does not present highly inter-species conserved regions, thus ensuring higher specificity. A LAMP primer set, named LAMP-Pfk13, was designed based on the alignment of consensus reference sequences of all human-infective *Plasmodium* species, as shown in Fig. 1B. These included zoonotic *Plasmodium* species such as *P. knowlesi* and *P. cynomolgy* which have recently jumped to infect human hosts [23, 35]. All sequences used for the alignment are provided in fig. S1. This primer set spans an amplicon size of 219 bp and consists of 6 primers targeting 8 different regions as shown in Fig. 1C: two outer primers F3 and B3, two loop primers LF and LB and two inner primers FIP and BIP. Primer sequences can be found in table S1.

**Fig. 1.**
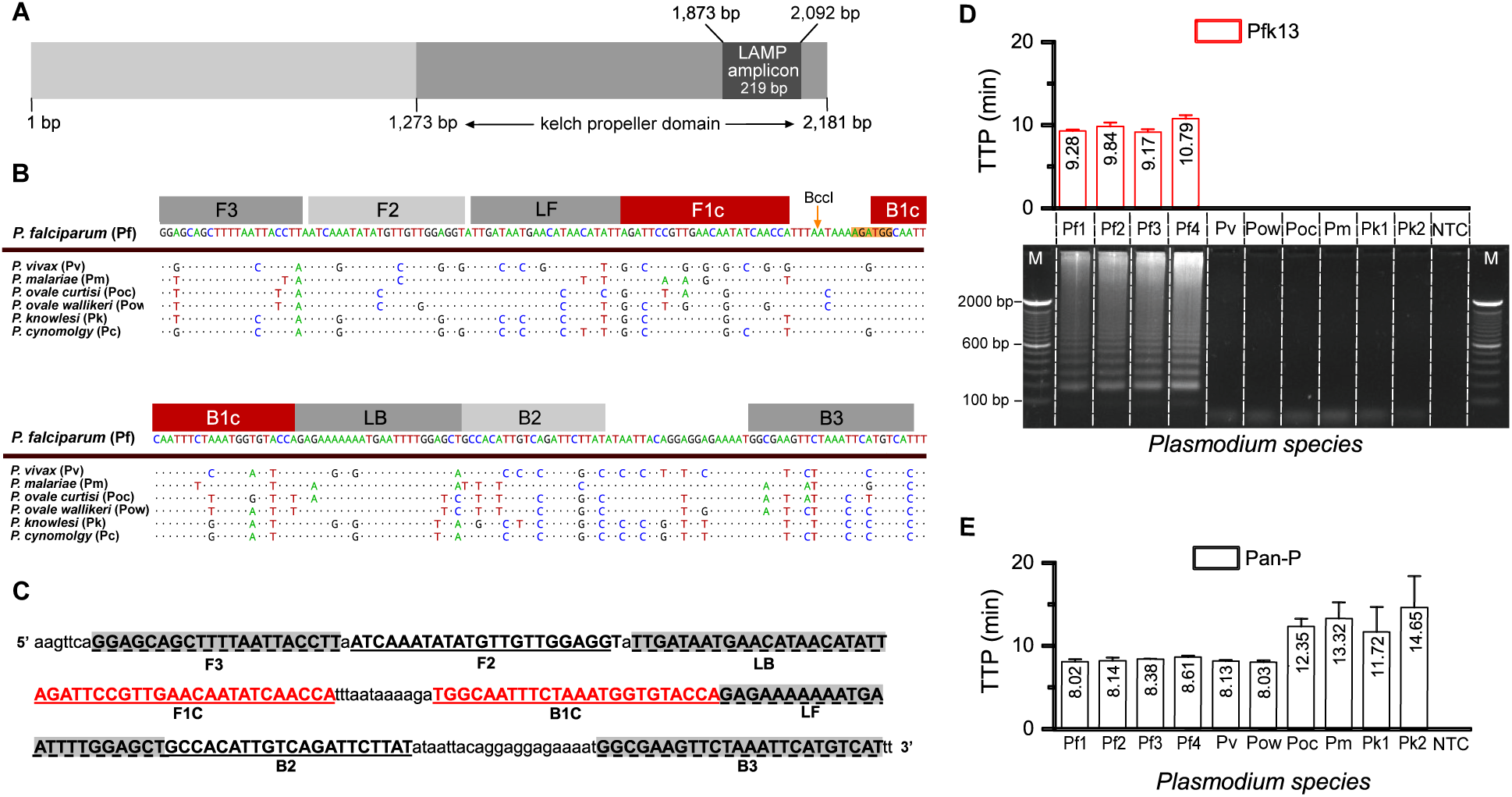
Sequence alignment, primer design and analytical specificity for the detection of *P. falciparum*. **(A)** Illustration of the gene *kelch 13* showing in black the region selected for primer design. **(B)** Alignment of all human-infective *Plasmodium* species with *P. falciparum* as reference sequence. Mismatches in the alignment are displayed as AGTC whereas matched nucleotides are represented as dots. Primers are illustrated on top of the alignment. **(C)** Primer location in LAMP amplicon (5’ to 3’ direction). Sequences of the primers can be found in table S1. **(D)** Results showing specific amplification of *P. falciparum* DNA samples from clinical isolates as well as gel electrophoresis of the amplified products to confirm specificity. Digested amplification products are shown in fig. S2. The restriction enzyme used for this experiment is BccI (# R0704S, New England BioLabs). The cutting point is illustrated with an arrow and the binding region is orange shadowed in (B). M denotes 100 bp DNA ladder (#10488058, Invitrogen). Samples are described in (B). **(E)** Pan-primer set [21] used as reference for the detection of all *Plasmodium* species. TTP values are displayed in minutes and labelled.

To illustrate the primer specificity, several human-infective *Plasmodium* species (*P. falciparum, P. malariae, P. vivax, P. ovale curtisi, P. ovale wallikeri* and *P. knowlesi)* were tested using a commercial LC96 qPCR instrument. Fig. 1D shows the detection of *P. falciparum* samples (Pf1 and Pf2) by including the time-to-positive (TTP) results obtained from the amplification reactions. The sample set included an additional two samples with mutations related to artemisinin-resistance (Pf3 denoting *P. falciparum* sample carrying the mutation *C580Y*, and Pf4 *P. falciparum* sample carrying the mutation *Y493H)*. To further confirm specificity, the same figure shows the bands of the LAMP amplified products obtained from agarose gel electrophoresis. Digested amplified products with the restriction enzyme BccI (#R0704S, New England Biolabs) are shown in fig. S2. These results demonstrate the absence of cross-reactivity across the *Plasmodium* species. In comparison, Fig. 1E shows the amplification responses when employing a Pan-Plasmodium primer set [21] which is able to detect all species. Overall, the, designed primer set is shown to be specific to the *P. falciparum* species.

Prior to this study, only one PCR primer set was reported targeting the gene *kelch 13* [25] yet without any information about the limit of the detection. To the best of our knowledge, this is the first isothermal assay targeting the gene *kelch 13* with high specificity for *P. falciparum* identification.

### 2.2 Analytical sensitivity of *P. falciparum* LAMP primer set

The novel LAMP assay, LAMP-PfK13, was tested with ten-fold serial dilutions of *P. falciparum* synthetic DNA ranging from 10^7^ to 10^0^ copies per reaction. Amplification curves from reactions carried-out in the LC96 qPCR instrument are included in the fig. S3. For quantification, the TTP values were obtained using the cycle-threshold metric, *C*_*t*_, set at 0.2 normalized fluorescence units by the LC96 instrument. Standard curves from these results are shown in Fig. 2A with high linearity (*R*^2^ = 0.993), a limit of detection of 10^0^ copies/reaction and maximum TTP of less than 20 minutes. Therefore, rapid quantification of unknown samples is possible with high reliability.

**Fig. 2.**
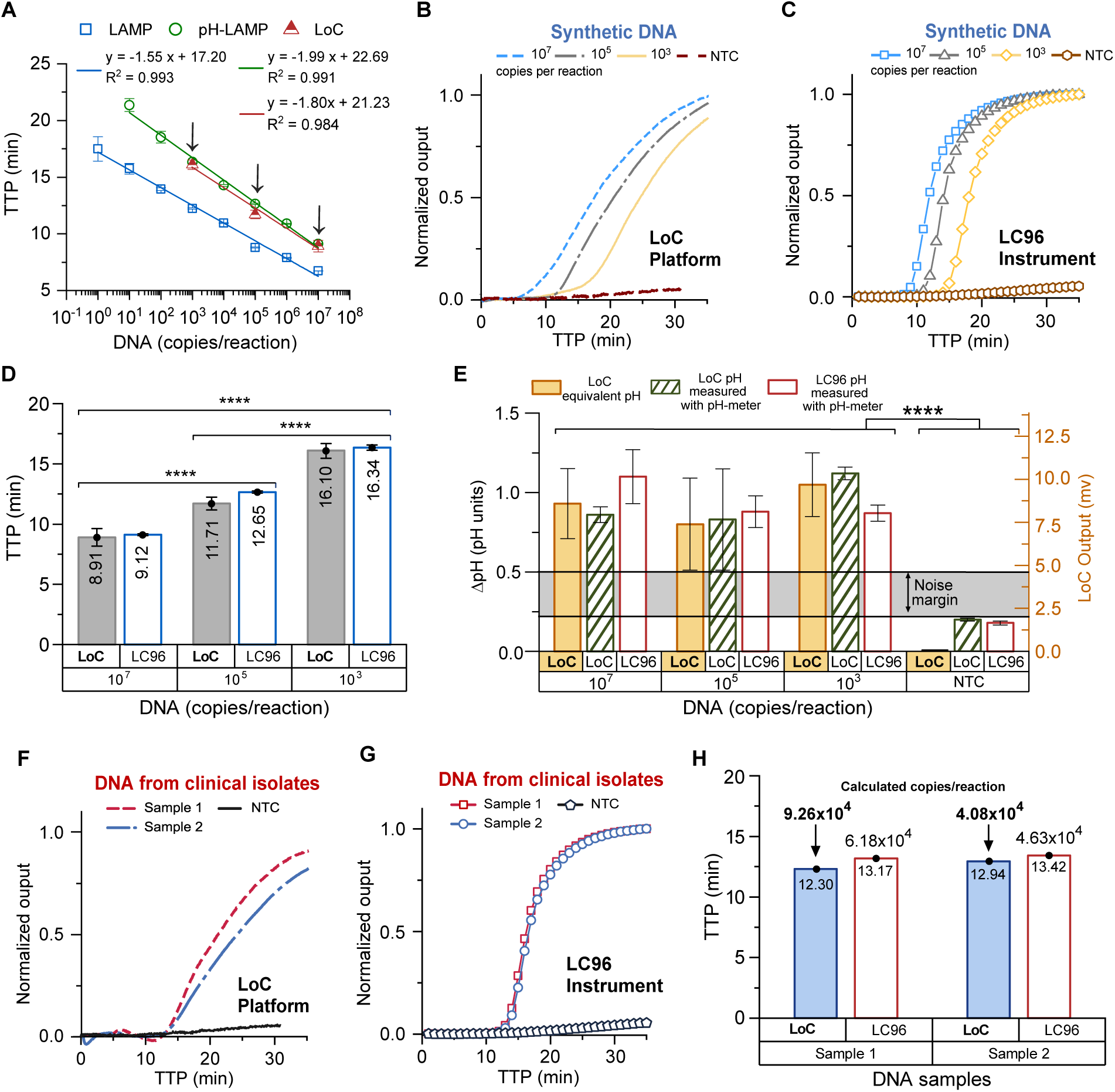
Summary of results of *P. falciparum* amplification with the LoC platform and a commercial qPCR instrument. **(A)** LAMP, pH-LAMP and LoC pH-LAMP standard curves obtained with the primer set LAMP-PfK13. Both pH-LAMP curves are almost perfectly correlated to the LAMP reference. **(B)** Amplification curves of synthetic DNA at different concentrations (10^7^, 10^5^ and 10^3^ copies per reaction) carried-out in the LoC platform. **(C)** Same samples carried out in LC96 qPCR instrument. **(D)** Bar plot comparing the TTP values obtained with the LoC platform and the LC96 qPCR instrument at the different concentrations of synthetic DNA. P-values produced from the Student’s t-test are shown as stars, with **** representing p-value< 0.0001. **(E)** Bar plot showing ΔpH measurements of the reaction solutions obtained before and after incubation at 63°C with the LoC (green bars) and the LC96 qPCR instrument (red bars). Furthermore, the LoC signal output (blue bars) in mV is shown (right-hand side Y-axis) and equivalent pH units according to the sensitivity described in Section 5 (left-hand side Y-axis). **(F)** Amplification curves of two *P. falciparum* DNA samples derived from clinical isolates carried-out in the LoC platform. **(G)** Same samples carried-out in the LC96 qPCR instrument. **(H)** Bar plot comparing the TTP values obtained with the LoC and LC96 instrument of the clinical isolates in (F) and (G). Also annotated are the equivalent concentrations estimated using the pH-LAMP standard curves in (A).

In order to transfer amplification chemistries on a Lab-on-Chip platform, they need to be compatible with the sensing capabilities of ISFET sensors. Since ISFETs are intrinsically pH sensors (mechanism for pH sensitivity elaborated in Section 5), the reaction mix of standard LAMP was modified to pH-LAMP by adjusting the buffering capabilities to allow for changes in pH to occur during DNA amplification. The corresponding amplification curves of pH-LAMP are shown in fig. S3 and standard curves in Fig. 2A. In this case, the limit of detection achieved was 10^1^ copies per reaction with an associated *R*^2^ = 0.991 and maximum TTP of less than 23 minutes.

Furthermore, to improve the robustness of extracting TTP in the case of sub-optimal amplification conditions, an alternative metric to *C*_*t*_, called *C*_*y*_, has been considered. This metric has been shown to be more reliable than *C*_*t*_ when considering samples of different efficiencies and is described by the intersection point between the time axis (or cycles) and the tangent of the inflection point of the amplification response [36]. For comparison, both *C*_*y*_ and *C*_*t*_ values have been derived from the fluorescent data obtained earlier with the LC96 instrument, and we show that they are perfectly correlated i.e. following a linear and monotonic relationship. The specific data considered are included in table S2. As a result, the *C*_*y*_ metric is also suitable for quantification purposes and has been used here when considering LoC data which do not rely on fluorescent measurements.

### 2.3 CMOS-based Lab-on-Chip detection and quantification of *P. falciparum*

Since the designed primer set enables specific, sensitive and rapid detection of *P. falciparum*, this section describes the results obtained by conducting amplification reactions on the Lab-on-Chip platform shown in Fig. 3A. The experimental setup includes a thermal controller, a microfluidic chamber for hosting the reaction mix and a sensing microchip comprising 4096 ISFET pixels. Cross-section of the ISFET fabricated in unmodified CMOS technology and the equivalent circuits are illustrated in Fig. 3A and b. Specific DNA amplification will modify the overall pH of the loaded solution causing a proportional electronic (voltage) change sensed by the ISFETs, illustrated in Fig. 3D. In the presence of a non-infected sample, amplification will not occur and the pH of the solution will remain the same, leading to a constant voltage signal. The temperature of the microchip is monitored through an electronic sensor designed as part of the microchip. More details on the methods and materials used, including a description of ISFET operation and data processing techniques are included in Section 5.

**Fig. 3.**
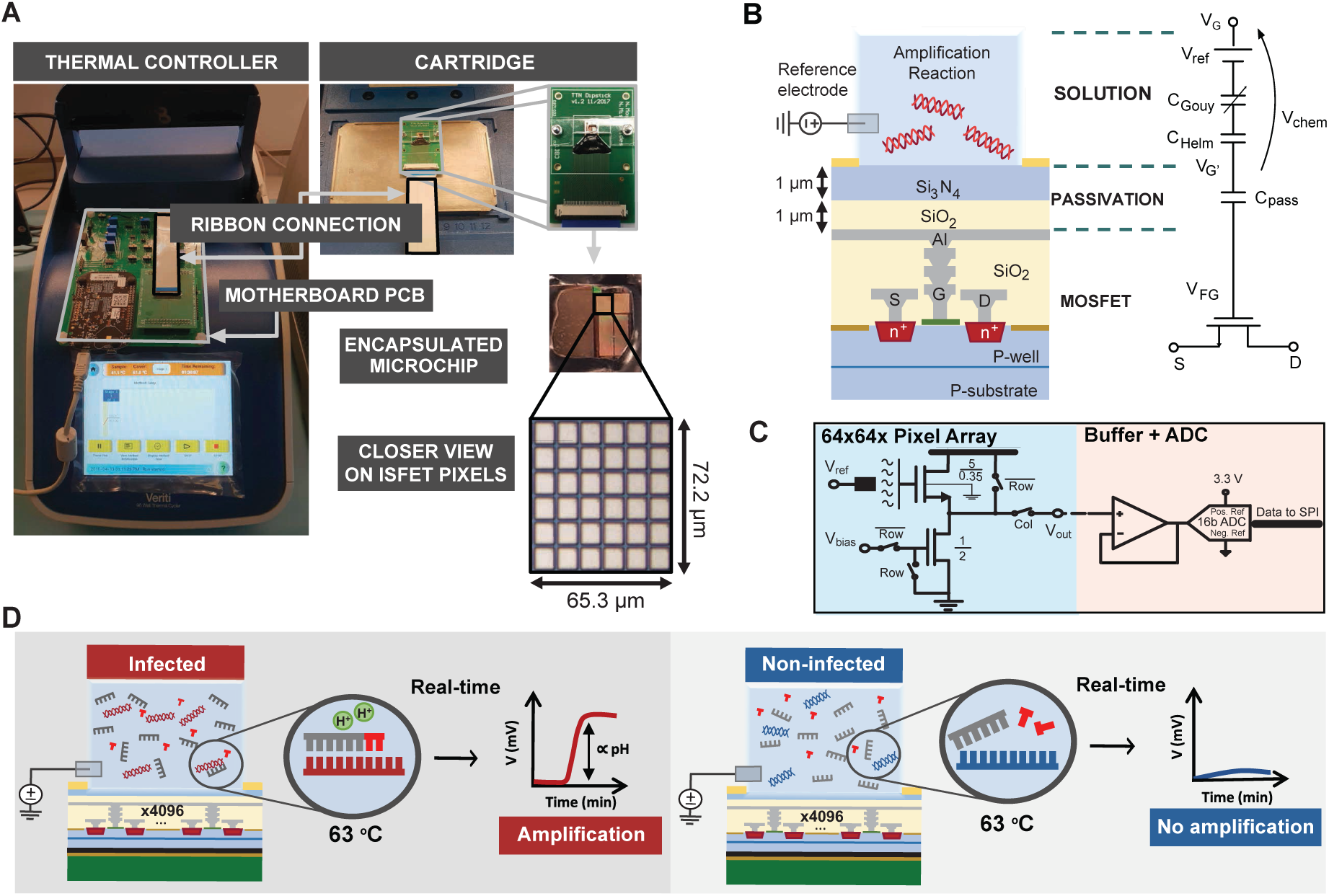
Experimental configurations using the Lab-on-Chip platform. **(A)** Setup showing the Lab-on-Chip (LoC) platform including a motherboard PCB that facilitates data readout, a cartridge PCB hosing the microchip and microfluidic chamber, a the microchip including an array of 4096 ISFET sensors and an external thermal controller. Furthermore, a microphotograph shows a 6×6 subset of the ISFET array. **(B)** Cross-section of an ISFET fabricated in unmodified CMOS technology and equivalent circuit macromodel. **(C)** Schematic of a pixel configuration implemented as a source follower configuration where changes in *V*_*out*_ reflect changes in pH [37]. **(D)** Cross-section illustration of the LoC platform showing the reaction interface. Amplification at 63°C only occurs if the specific target is present in the loaded sample. Results are displayed in real-time.

Several concentrations of *P. falciparum* synthetic DNA (10^7^, 10^5^ and 10^3^ copies per reaction) were tested in the LoC platform to evaluate its capability for nucleic acid amplification and quantification. Amplification responses from carrying-out pH-LAMP, in the form of pH-to-voltage signals, were post-processed in software to compensate for known ISFET non-idealities such as sensor drift and thus obtain (i) average and normalised amplification curves and (ii) TTP values using the *C*_*y*_ method[36]. The standard curve built with LoC data and used for quantification is shown in Fig. 2A, with the LoC amplification curves in Fig. 2B. In comparison, Fig. 2C shows the respective responses from the LC96 instrument used as reference. The TTP values obtained across the various DNA concentrations from both the LoC and LC96 are illustrated and compared in Fig. 2D. Statistical analysis of these results using Student’s t-test to compare means as well as correlation test to compare trends, report a p-value <0.0001 and a correlation coefficient of 0.99 (more details in table S3). These metrics indicate that there exists an insignificant probability of unequal concentrations producing similar TTP values and that the trends across concentrations are almost perfectly dependent. Overall, the robustness of the LoC against a commercial instrument is demonstrated for DNA quantification which is consistent with the standard curves presented earlier in Fig. 2A.

Deviations between the TTP values obtained with the LoC platform and the LC96 instrument could be attributed to several factors such as (i) intrinsic error among technical replicates, (ii) reaction conditions variability such as different material of the reaction chamber, (iii) chip non-idealities such as trapped charge or variations in the sensing *Si*_3_*N*_4_ layer, and (iv) different nature of the detected signal (pH-vs-fluorescence). Specifically, fluorescence measurements are obtained as the intercalating dye is being incorporated in newly generated amplicons [38–40]. On the contrary, in the LoC platform the chemical input in the form of H^+^ concentration (*pH* =−*log*[*H*^+^]) is converted to an electrical output. Multiple amplification events take place simultaneously in LAMP, and therefore, describing the kinetics of this assay is not trivial as well as establishing a relationship between fluorescence and change in pH [41–43]. Nevertheless, the high degree of correlation recorded between the TTP values obtained using both types of responses implies that both can be used as indicators of the amplification state, something that was previously argued in [32].

To further confirm LoC amplification, the pH of the solution was measured before and after incubation at 63°C. Specifically, Fig. 2E shows the ΔpH obtained from running reactions of the same concentration in the LC96 instrument as well as in the LoC, with the solution pH measured using a commercial (Sentron SI600) pH meter.pH measurements and DNa quantification values can be found in table S4 and S5. In addition, the figure includes the equivalent pH obtained from the output of the ISFET sensors with the previously determined sensitivity of 9.23 mV/pH. A small pH change occurs for non-template control samples, never-theless a wide margin of error was obtained which can be used to set a pH threshold indicating amplification through the recorded pH change.

To demonstrate the quantification capabilities, two *P. falciparum* gDNA samples derived from clinical isolates were amplified and quantified using both the LoC and reference LC96 instrument. Amplification curves and TTP values obtained from both reactions are shown in Fig. 2F-H. Using the previously derived standard curves, the estimated LoC initial concentration values are within <4% from the reference demonstrating the capability of the LoC to quantify clinical isolates accurately.^1^

### 2.4 LoC detection of *P. falciparum* drug-resistant malaria

The most common mutation indicating the presence of drug-resistant parasites is the *C580Y* single nucleotide polymorphism (SNP). The LAMP-based method described in Malpartida-Cardenas *et al.* [44] reported the discrimination of wild-type (WT) from mutant (MT) alleles by robustly preventing or delaying unspecific amplification, as illustrated in 4A. Two reactions are tested, one targeting the presence of the WT allele (WT reaction) and another one targeting the presence of the MT allele (MT reaction). Primer sequences can be found in table S6. In this case, this method was adapted to pH-LAMP in order to be transferred onto the Lab-on-Chip platform.

*P. falciparum* DNA samples from clinical isolates known to be WT or MT were tested with the LoC platform and with the LC96 qPCR instrument as control. Results are presented in Fig. 4B, showing the TTP values obtained with each reaction. Overall, SNP discrimination is possible due to delayed amplification of unspecific targets leading to a difference in TTP values (Δ TTP). As a result, the capability of the LoC platform to discriminate alleles is illustrated in the same way as the commercial LC96 qPCR instrument.

**Fig. 4.**
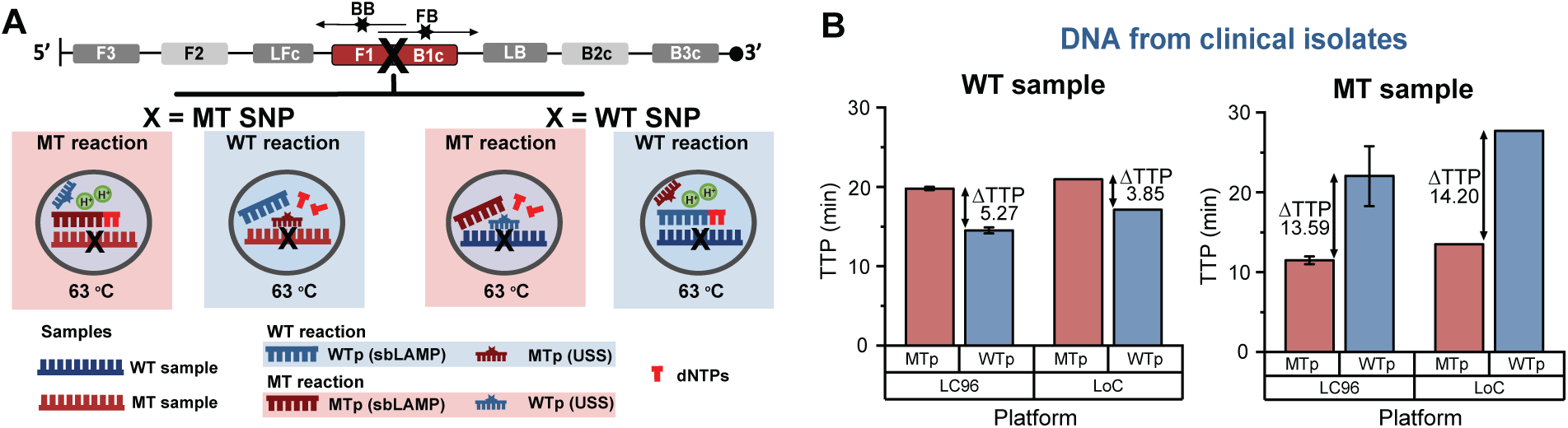
Lab-on-Chip detection of *C580Y* SNP associated to *P. falciparum* artemisinin-resistance. **(A)** Workflow representation of the USS-sbLAMP method for allele-specific detection [44]. Specific reactions amplify in isothermal conditions whereas unspecific reactions are prevented or delayed. **(B)** Comparison of results obtained with the LoC platform and the LC96 qPCR instrument. TTP values are displayed, with ΔTTP values annotated, indicating that specific reactions always occur earlier.

## 3 Discussion

Our results demonstrate that the developed LAMP assays paired to the LoC platform are well-suited for clinical applications with significant potential towards point-of-care implementations. This is achieved by tailoring amplification chemistries to be compatible with electronic detection using an ISFET-based platform and a Lab-on-Chip approach. We demonstrate the potential of such an implementation for the detection of malaria-causing *P. falciparum* using a novel and specific primer set as well as specific identification of drug-resistant mutations. Furthermore, we show for the first time *P. falciparum* DNA quantification on a LoC platform with high accuracy and robustness comparable with a commercial benchtop instrument.

Compared to other rapid diagnostic tests for pathogen detection, we opted for nucleic-acid-based due to their high sensitivity and specificity. Even though PCR is considered the gold standard, in this case isothermal amplification using LAMP avoids the need for complex temperature control, is more specific and rapid and is therefore more suitable for point-of-care applications. Furthermore, no pre-treatments at different temperatures are needed for optimal DNA amplification, as opposed to the conventional SDA method[45]. Results obtained with the LAMP assay targeting the gene *kelch 13* in *P. falciparum* show high specificity without cross-reactivity across other *Plasmodium* species and high sensitivity with a limit of detection of 1 copy per reaction. In contrast with prior works [19, 21, 27], the speed of detection shown here is less than 20 min. In addition, modifying the reaction chemistries to pH-LAMP for electrochemical compatibility is achieved with almost identical behaviour, indicating the feasibility of carrying-out the reactions in an ISFET-based platform.

The developed LoC platform leverages on ISFET-based detection to facilitate fully-electronic chemical sensing and direct electrochemical signal transduction [37]. Sensor fabrication in CMOS technology achieves sensor miniaturisation, low cost manufacturing, scalability and sensitivity which are necessary for a point-of-care implementation. Furthermore, detection takes place with sensors fabricated in unmodified CMOS technology without the use of probes, electrodes or prior surface treatment. LoC detection has been demonstrated for DNA amplification reactions with results matching those from a commercial benchtop instrument and allowing for building the first LoC standard curve. The standard curve was validated with unknown DNA samples from clinical isolates with high accuracy. In addition, SNP discrimination on the LoC platform was achieved with DNA samples harbouring the *C580Y* mutation derived from clinical isolates.

Nevertheless, to fully realize a PoC platform for *P. falciparum* and drug-resistant malaria detection, several issues need to be optimised further.

Firstly, the current workflow still relies on the use of extracted DNA rather than samples directly obtained from patients (e.g. from bodily fluids most typically blood samples). Consequently, the introduction of a sample preparation step is necessary in order for such an approach to be fully deployed as Point-of-Care sample-to-result platform. Nevertheless, isothermal techniques and specifically LAMP which was employed here, have been shown to be more robust to typical blood inhibitors compared to PCR [27] which can be leveraged to simplify such a sample preparation process. Secondly, from an electronic aspect, embedded temperature control is key for achieving full portability since temperature in the LoC platform presented here is controlled using an external instrument. That extends to enclosing all electronic functions into portable packaging with embedded temperature control, isolating the microchip and microfluidic modules in the form of a disposable cartridge from the base unit responsible for data readout and processing. Towards this direction, we have already designed a reusable motherboard and a disposable cartridge module. The cartridge only includes the sensing microchip, microfluidic setup and amplification reagents while leveraging on CMOS technology to ensure mass manufacture of sensors and thus low-cost, suitable for point-of-care applications.

In summary, the proposed LoC platform in combination with novel isothermal chemistries for *P. falciparum* specific DNA detection have demonstrated performance comparable to a commercial benchtop instrument. Overall, we have demonstrated DNA amplification and quantification, as well as SNP discrimination. We anticipate that such a LoC platform holds significant potential for full deployment in resource-limited settings, towards the rapid diagnosis of infectious diseases and reduction of antimicrobial resistance.

## 4 Molecular Methods

### 4.1 LAMP primer design specific to *Plasmodium* falciparum

A LAMP primer set for the detection of the gene *Kelch 13* of *P. falciparum*, named LAMP-PfK13, was designed based on the alignment of consensus reference genomic sequences from all human-infective *Plasmodium* species and some zoonotic *Plasmodium* species including *P. knowlesi* and *P. cynomolgi*. Sequences were retrieved from *Plasmodium* Genomic Resource (PlasmoDB) [46] and aligned using the MUSCLE algorithm [47] in Geneious 10.0.5 software [48]. Gene IDs of the sequences used for the alignment can be found in fig. S1. The LAMP primer set LAMP-PfK13 was designed using Primer Explorer V5^2^ and optimized manually to ensure specificity and to locate the loop primers (LB and LF). Sequences of the primers can be found in table S1.

### 4.2 Reaction conditions

Each LAMP reaction contained the following: 1.5 µL of 10× isothermal buffer (New England Biolabs)^3^, 0.9 µL of MgSO4 (100 mM stock), 2.1 µL of dNTPs (10 mM stock), 0.375 µL of BSA (20 mg/mL stock), 2.4 µL of Betaine (5M stock), 0.375 µL of SYTO 9 Green (20 µM stock), 0.6 µL of Bst 2.0 DNA polymerase (8,000 U/mL stock), 3 µL of different concentrations of synthetic DNA or gDNA, 1.5 µL of 10×LAMP primer mixture (20 µM of BIP/FIP, 10 µM of LF/LB, and 2.5 µM B3/F3) and enough nuclease-free water (ThermoFisher Scientific) to bring the volume to 15 µL. Reactions were performed at 63°C for 35-40 min. One melting cycle was performed at 95°C for 10 s, 65°C for 60 s and 97°C for 1 s for validation of the specificity of the products. Experiments were carried out twice and each condition was run in triplicates (5 µL each reaction) loading the reactions into LightCycler 480 Multiwell Plates 96 (Roche Diagnostics) utilising a LightCycler 96 Real-Time PCR System (Roche Diagnostics). In the case of pH-LAMP, the buffering conditions were modified such that pH changes could be measured. Each reaction contained the following: 3.0 µL of 10× isothermal customized buffer (pH 8.5 - 9), 1.8 µL of MgSO4 (100 mM stock), 4.2 µL of dNTPs (10 mM stock), 1.8 µL of BSA (20 mg/mL stock), 4.8 µL of Betaine (5 M stock), 1.88 µL of Bst 2.0 DNA polymerase (8,000 U/mL stock), 3 µL of different concentrations of synthetic DNA or gDNA, 0.75 µL of NaOH (0.2 M stock), 3 µL of 10 ×LAMP primer mixture, 0.75 µL of SYTO 9 Green (20 µM stock) only for qPCR experiments, and enough nuclease-free water (ThermoFisher Scientific) to bring the volume to 30 µL. Experiments were conducted twice and each condition was ran in duplicates or triplicates (10 µL each). Reactions tested on chip were carried-out in duplicates (13 µL each). Synthetic DNA sequences used in this work were purchased from Integrated DNA Technologies (The Netherlands) and resuspended in TE buffer to 100 µM stock solutions.

### 4.3 Samples and DNA extraction methods

A gBlock Gene fragment containing the *kelch 13* region of interest was purchased from Integrated DNA Technologies and re-suspended in TE buffer to 5 ng/µL stock solution, stored at −20°C. *Plasmodium* genomic DNA (gDNA) of all human-infective *Plasmodium* species (*P. falciparum, P. ovale curtisi, P. ovale wallikeri, P. vivax, P. malariae* and *P. knowlesi)* were tested.^4^ *P. falciparum* genomic gDNA samples containing the *Y493H* mutation (ANL8G), and the *C580Y* mutation (ANL5G) in this gene were isolated using the PureLink Genomic DNA Mini Kit (ThermoFisher Scientific) from Cambodian isolates.

### 4.4 Analytical sensitivity of *P. falciparum* specific primer set

Sensitivity of the LAMP-PfK13 primer set was evaluated using ten-fold serial dilutions of synthetic DNA ranging from 1×10^7^ to 1×10^0^ copies per reaction with LAMP and pH-LAMP. For each method, a standard curve was obtained by plotting the time to positive (TTP) against copies per reaction accompanied by the standard deviations.

### 4.5 Cross-Reactivity and detection of clinical isolates

Ten DNA samples derived from clinical isolates were tested with the LAMP-PfK13 primer set to prove the absence of cross-reactivity with any other human-infective *Plasmodium* species. Samples (gDNA) included: *P. ovale curtisi, P. ovale wallikeri, P. vivax, P. malariae*, two *P. knowlesi* samples and four *P. falciparum* samples with one harbouring the mutant allele *580Y*, one more harbouring the mutant allele *493H* and two wild type samples.

### 4.6 SNP discrimination on chip

In the case of specific SNP detection, mutant (MT) and wild-type (WT) reactions were performed as described in Malpartida-Cardenas *et al.* [44]. Sequences of primers can be found in table S6. These reactions consisted of SNP-based LAMP (sbLAMP) primers targeting the specific allele for amplification (i.e MT allele), and unmodified self-stabilising (USS) competitive primers targeting the other allele (i.e WT allele) to robustly delay or prevent unspecific amplification. Each USS-sbLAMP reaction followed the usual LAMP reaction protocol with the inclusion of 10× sbLAMP primer mixture (20 µM of sbBIP/sbFIP, 10 µM of LF/LB, 2.5 µM B3/F3 and 3 or 4 µM of *FB/BB*), rather than 10× LAMP. USS (Integrated DNA Technologies) were re-suspended in TE buffer to 400 µM. Reactions were carried-out at 63°C for 35 min. This method was modified to be pH-sensitive such that it could be utilised in combination with the LoC platform as described in 4.2.

### 4.7 Gel Electrophoresis

Agarose gels were prepared at 1.5% w/v with TBE 1× buffer and SYBR Safe DNA Gel Stain 1000×. LAMP products were mixed with loading dye (#B7024S, New England BioLabs) at 6x concentration (2µL) prior to loading into pre-cast wells in the gel. As reference, 100 bp DNA ladder (#10488058, Invitrogen) was loaded. Power supply was set at 1-5V/cm to run the gel for 1h. Stained DNA was visualized under UV light with UV BioSpectrum Imaging System instrument (Ultra-Violet Products Ltd.).

### 4.8 Statistical Analysis

Data is presented as mean TTP ± standard deviation, in minutes; p-values were calculated by Student’s heteroscedastic t-test, with a two-sided distribution. Statistically significant difference was considered as: *p-value < 0.05, **p-value < 0.01, ***p-value < 0.001 and ****p-value < 0.0001. Correlation coefficients were calculated using Eq. 1 in table S2.

## 5 CMOS-based Chemical Sensing

### 5.1 ISFET-based Sensing

Detecting pH changes in an electrochemical manner is facilitated using ISFETs fabricated in unmodified CMOS technology [49]. ISFETs are designed the same way as MOSFETs with the gate extended to the top metal layer using a floating metal stack. With this method, the gate is biased using a reference electrode (typically Ag/AgCl) immersed in a solution. Sensing takes place through the passivation layer or the deposition of a suitably selective membrane which exhibits site binding and develops a double layer capacitance when exposed to an aqueous solution. The accumulation of protons due to the combined effect of the electrode biasing and the hydrogen ion concentration in the solution is capacitively coupled to the floating gate and modulates the gate potential of the underlying device. When biased with a stable reference electrode voltage, variations at the floating gate potential can be attributed to changes in ion concentration in the solution therefore a pH dependence is observed.

The equivalent circuit macromodel of an ISFET in unmodified CMOS technology is shown in Fig.3b with the chemical dependence described by a term called *V*_*chem*_ [49] given by:

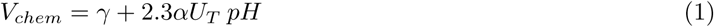

where *γ* describes all the constant terms not related to pH, *α* is a dimensionless sensitivity parameter and *U*_*T*_ is the thermal voltage.

Owing to their compatibility with modern electronic processes and the economies of scale of silicon, ISFET-based sensing has the potential for miniaturized, mass-fabricated and low-cost solutions. As a result, microchips with integrated ISFET sensing provide an attractive silicon substrate for LoC applications. However, sensor non-idealities exist which impose additional challenges on chemical sensing in unmodified CMOS [50]. Firstly, trapped charge is typically left during fabrication at the floating gate and manifests as a random offset across sensors. This can be compensated by introducing sensor redundancy to reduce the susceptibility to high offsets and ensuring a large dynamic range in which sensors operate linearly irrespective of offsets. Secondly, the sensing membrane undergoes hydration after being exposed to an aqueous solution which manifests as a slow change on the output signal, typically referred to as drift. Drift is typically slower than pH changes due to DNA reaction dynamics and generic compensation methods have typically revolved around derivative-based methods [51]. In this case, a compensation method is shown to decouple the local changes in pH from the background drift.

### 5.2 ISFET sensing array

An array of 64 × 64 ISFET pixels of identical geometry has been fabricated in the AMS 0.35µm process using silicon nitride (*Si*_3_*N*_4_) as the passivation layer. Each pixel contains an ISFET configured in a source follower topology as shown in Fig. 3A with the pixel output buffered and sampled using an external 16-b ADC. This way changes in pH are linearly converted into changes in *V*_*out*_ for active pixels in the linear input range i.e. excluding extreme cases of trapped charge. Each pixel spans approximately 96 *µm*^2^ of silicon area with the total array of 4096 sensors spanning 0.56 *mm*^2^. The intrinsic pH sensitivity of the *Si*_3_*N*_4_ layer has been measured to be 18 *mV/pH* with the final sensitivity of *V*_*out*_ to pH after capacitive attenuation measured to be 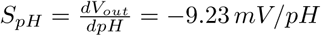. A detailed description of the circuit design, instrumentation pipeline and method to derive the pH sensitivity is provided in Miscourides & Georgiou [37].

### 5.3 Experimental Setup

To facilitate data readout and communication to a PC, a printed-circuit-board (PCB) was designed to host a microcontroller serving as an intermediary node between the microchip and a PC. Furthermore, a separate cartridge PCB was designed to host the microchip which is connected with a ribbon cable to the main PCB and communication takes place via the serial peripheral interface. A Matlab-based graphical user interface is used to control all operations and provide real-time data recording and visualisation at 0.3 fps.

Moving towards a LoC platform to carry out on-chip DNA amplification detection, a microfluidic reaction chamber was laser cut from a 3 mm acrylic sheet and assembled on top of the CMOS microchip as shown in Fig. 3A. In addition, a Ag/AgCl reference electrode was inserted in the chamber for sensor biasing, which was obtained by chloridation of a 0.03 mm diameter Ag wire in 1M KCl.

To carry-out DNA amplification reactions, the microfluidic chamber was filled with 13 µL of pH-LAMP or pH-USS-sbLAMP reagents and sealed with PCR tape to avoid evaporation and contamination of the amplified products. Subsequently, the cartridge PCB was placed on top of a thermal cycler (Veriti Thermal Cycler, Applied Biosystems) used as a temperature controller to keep the solution at 63°C for isothermal amplification. After 35-40 min, the solution was recovered for further analysis. Measurements of pH with a commercial pH meter line (Sentron SI600) and DNA quantification with a fluorometer (Qubit 3.0, Thermo Fisher Scientific) were obtained from the recovered solution to check whether amplification occurred as well as the corresponding pH change.

Furthermore, Figs. 5a-c show the response of the temperature sensor include on-chip to monitor the temperature during DNA amplification reactions. The sensor is based on a typical PTAT circuit configuration with a linear response.

**Fig. 5.**
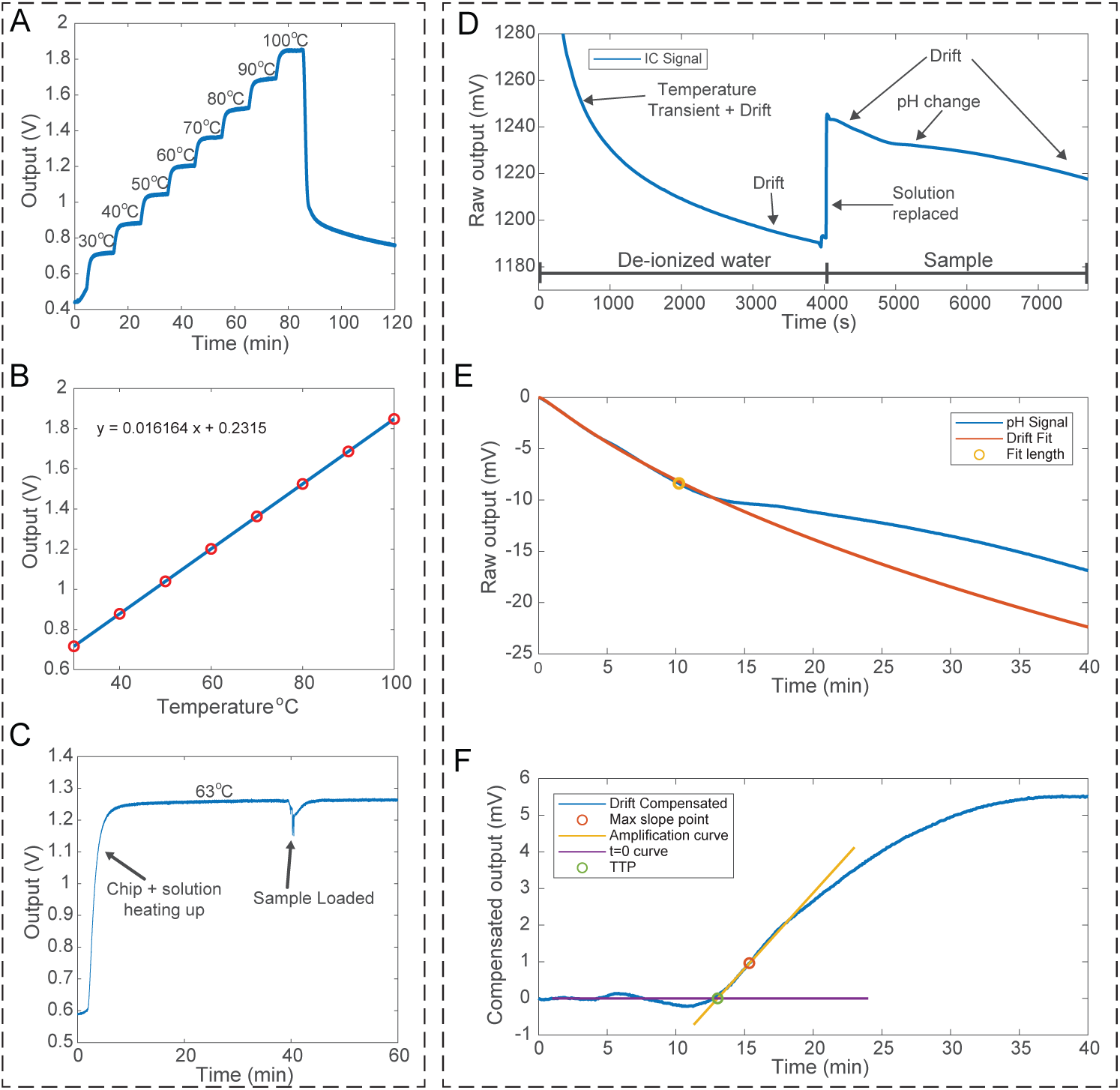
Electronic Sensors and Methods. **(A)** Temporal characterisation of the temperature sensor on-chip at steps of 10°C. **(B)** Temperature sensor linearity showing a sensitivity of 16.2 mV/°C. **(C)** Temperature profile from the on-chip sensor during a typical reaction. **(D)** Typical response obtained during pH-LAMP carried-out in the LoC platform. The trace shows raw data recorded as the average of active pixels in the ISFET array. At first, the chip heats up with de-ionized water loaded in the chamber and a temperature transient is observed. Subsequently, the slow monotonic change shown corresponds to drift due to hydration of the sensing material. After the temperature has settled, the solution is replaced with a positive sample which induces a pH change. **(E)** Signal recorded from the ISFET sensors after the sample has been loaded. The stretched-exponential drift model is adopted whose parameters are fitted using temporal data from the first few (¡10) minutes of the reaction. Subsequently, drift is extrapolated until the end of the amplification reaction and is assumed as the background signal. **(F)** Drift-compensated signal with the TTP obtained by finding the point where the amplification curve crosses y=0. The amplification curve is obtained as the straight line at the point of maximum slope of the drift-compensated signal.

### 5.4 Mechanism of pH-LAMP detection using ISFETs

During nucleic acid amplification, nucleotides are incorporated by action of a polymerase resulting in the release of a proton (H^+^) as described in Eq. 2. The release of protons into the solution induces a change in pH that ISFETs can transduce into an electrical output, thus correlating a change in pH to a change in voltage [32].

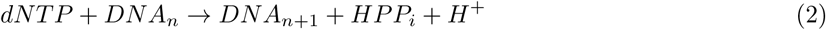

Additionally, this change is regulated by the buffer capacity (*β*_*int*_) of the solution. Overall, the change in pH is given by:

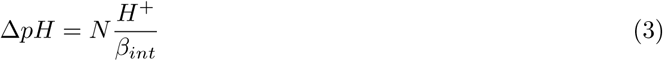

where *N* denotes the total number of nucleotides incorporated during amplification. Consequently, the pH change is proportional to the total number of nucleotides inserted and inversely proportional to the buffering capacity of the solution. As a result, this opens up the possibility of electronic sensing, whereby ISFETs can be used to track the pH change during nucleic acid amplification. The expected modification in ISFETs is described by modifying Eq. 1 to:

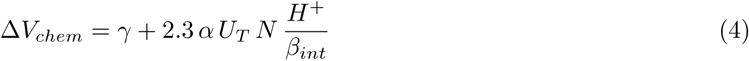

This equation can be used to describe the amplification curve which follows an exponential profile. At the early stages of the reaction, the amount of protons generated is not enough to overcome the buffering capacity of the solution and could also be very small compared to the sensitivity of the sensing layer. Eventually, the amount of protons accumulated is enough for showing a detectable (exponential) signal. Subsequently, as the amount of reagents is exhausted, the reaction profile enters the saturation stage and reaches a plateau.

### 5.5 Data analysis

During an amplification reaction, the pH signal is obtained by taking the average of the active sensors that are exposed to the solution. The signal includes drift due to the hydration of *Si*_3_*N*_4_ used for sensing which manifests as a slow, monotonic change on the output [51]. The underlying cause is the diffusion of ions from the solution to charge-trapping sites in the nitride that exist due to its amorphous structure. This phenomenon can be modelled as a dispersive transport process following a stretched-exponential response (exp[-*t*^*β*^]). Consequently, the first few minutes (< 10) of the amplification reaction are used to sample drift and derive an analytical equation that fits the drift observed. As a result, the pH signal can be decoupled from the expected drift and the pH change due to DNA amplification can be obtained. To ensure accurate modelling of drift, the compensation method is applied locally around the exponential amplification point and is not considered valid at large extrapolated values of the drift model. Responses before and after drift compensation are shown in Fig. 5D-F.

Furthermore, after obtaining the drift-compensated signal which captures the pH change due to amplification, the time-to-positive (TTP) metric is obtained. The *C*_*y*_ method is adopted to derive the TTP as the time when the linear extrapolation at the inflection point intersects the background signal[43].

The combination of these two methods is outlined below and illustrated graphically in Fig. 5. Bold notation (e.g. **v**) is used to indicate vectors or time series.

1. The pH signal **p** obtained from an amplification reaction is normalized by removing the DC component (background).
2. During the first few minutes (< 10) after the sample is loaded, sensor drift is modelled using 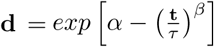 where the scalar parameters *α, τ* and *β* are estimated.
3. **d** is extrapolated until the completion of the amplification reaction.
4. The drift-compensated response is obtained using **p** − **d**.
5. The inflection point is determined as the point of maximum derivative of **p**−**d**. A linear response is fitted around the maximum derivative point (amplification curve).
6. TTP is defined as the time when the amplification curve crosses y=0.

Results obtained with the LoC platform prior to any processing steps are shown in fig. S4.

## Data and materials availability

All data are present in the paper and/or in the Supplementary Materials. Additional data related to this paper may be requested from the authors.

## Footnotes

1 The percentage change was calculated by referring the ion concentration values (i.e. 10^*x*^) to a linear axis.

2 Eiken Chemical Co. Ltd., Tokyo, Japan, http://primerexplorer.jp/lampv5e/index.html

3 Catalogue number B0537S

4 These samples were kindly provided by Prof. Colin Sutherland from The London School of Hygiene and Tropical Medicine.

